# ORF48 is required for optimal lytic replication of Kaposi’s Sarcoma-Associated Herpesvirus

**DOI:** 10.1101/2024.02.29.582672

**Authors:** Beatriz H S Veronese, Amy Nguyen, Khushil Patel, Kimberly Paulsen, Zhe Ma

## Abstract

Kaposi’s sarcoma-associated herpesvirus (KSHV) establishes persistent infection in the host by encoding a vast network of proteins that aid immune evasion. One of these targeted innate immunity pathways is the cGAS-STING pathway, which inhibits the reactivation of KSHV from latency. Previously, we identified multiple cGAS/STING inhibitors encoded by KSHV, suggesting that the counteractions of this pathway by viral proteins are critical for maintaining a successful KSHV life cycle. However, the detailed mechanisms of how these viral proteins block innate immunity and facilitate KSHV lytic replication remain largely unknown. In this study, we report that ORF48, a previously identified negative regulator of the cGAS/STING pathway, is required for optimal KSHV lytic replication. We used both siRNA and deletion-based systems to evaluate the importance of intact ORF48 in the KSHV lytic cycle. In both systems, loss of ORF48 resulted in defects in lytic gene transcription, lytic protein expression, viral genome replication and infectious virion production. ORF48 genome deletion caused more robust and global repression of the KSHV transcriptome, possibly due to the disruption of RTA promoter activity. Mechanistically, overexpressed ORF48 was found to interact with endogenous STING in HEK293 cells. Compared with the control cell line, HUVEC cells stably expressing ORF48 exhibited repressed STING-dependent innate immune signaling upon ISD or diABZI treatment. However, the loss of ORF48 in our iSLK-based lytic system failed to induce IFNβ production, suggesting a redundant role of ORF48 on STING signaling during the KSHV lytic phase. Thus, ORF48 is required for optimal KSHV lytic replication through additional mechanisms that need to be further explored.

**Author Summary:** Kaposi sarcoma-associated herpesvirus (KSHV) causes persistent infection in a host that leads to two deadly cancers, Kaposi Sarcoma and Primary Effusion Lymphoma, especially in immunocompromised people. Unfortunately, there is no vaccine or viral-specific treatment for KSHV-related diseases, due to our limited knowledge of detailed immune evasion strategies by KSHV. KSHV blocks multiple immune pathways to maintain its lifelong infection, one of which is the DNA-sensing cGAS-STING pathway. Here, we reported that ORF48, a KSHV-encoded STING inhibitor is required for optimal KSHV lytic reactivation and viral production. A successful KSHV infection requires both intact ORF48 DNA and mRNA at different stages of its lytic life cycle. Further study reveals that ORF48 binds to STING and blocks STING-dependent innate immunity, and additional mechanisms may contribute to its role in lytic replication. Our findings provide insight into viral immune evasion strategies, which would contribute to a better understanding of all viral diseases.

## Introduction

Kaposi’s sarcoma-associated herpesvirus (KSHV) or human herpesvirus 8 (HHV8) is the etiological agent of multiple human malignancies, such as Kaposi sarcoma (KS), multicentric Castleman’s disease (MCD), primary effusion lymphoma (PEL), and KSHV-inflammatory cytokine syndrome(1–5). KSHV has two infection phases: latency and lytic replication(6). Latently infected cells express a reduced number of viral genes and no infectious virions are generated during this phase(7). On the contrary, the lytic cycle is characterized by the transcription of the entire KSHV genome and the production of infectious virion particles(8), thereby increasing the risk of immune detection of the virus by the host(9). KSHV lytic proteins must therefore exert immunomodulatory functions to enable persistent infection, many of which are yet to be explored. Of the more than ninety KSHV open reading frames (ORFs) that have been identified, several have been shown to contribute to immune evasion and facilitate the lifelong infections of KSHV(10–18).

KSHV blocks multiple immune pathways to maintain its persistent infection, one of which is the DNA-sensing cGAS-STING pathway(10,14,15,19). cGAS (cyclic GMP-AMP synthase) senses cytosolic DNA originating from pathogen infection or genome instability(20). It then catalyzes the formation of the second messenger cGAMP, which binds to and activates ER-located STING (stimulator of interferon genes, also known as MITA, ERIS, MPYS)(20–24). STING recruits TBK1 (TANK-binding kinase 1) and gets phosphorylated (25). IRF3 (Interferon regulatory factor 3) is then recruited to this complex and gets phosphorylated by TBK1(25). Lastly, phosphorylated IRF3 translocates to the nucleus and triggers the production of type I interferons (IFNs), a critical cytokine protecting hosts against viral infection(26). Loss of cGAS or STING in reactivated KSHV-harboring iSLK.219 cells resulted in attenuated IFNβ production throughout the lytic stage of KSHV, and led to significantly stronger viral lytic gene transcription, lytic protein expression, and infectious virions(10). Consistently, activating STING by cGAMP exhibits the opposite effect(15).

Further studies revealed multiple viral proteins and mechanisms that silence the cGAS/STING-based innate immunity. For instance, KSHV ORF52 was found to inhibit cGAS enzymatic activity, and therefore attenuate sufficient DNA-sensing by cGAS(14). KSHV viral interferon regulatory factor 1 (vIRF1) binds to STING and sequesters STING from being sufficiently phosphorylated by TBK1(10). In addition, a truncated LANA interacts with cGAS and negatively regulates the cGAS/STING-dependent type I interferon production(15). In addition, many host negative regulators of STING are hijacked by KSHV to facilitate its lytic replication, such as NLRX1 and PPM1G(27,28). Thus, KSHV needs to keep the cGAS-STING signaling repressed during its lytic cycle. Further characterization of other viral candidates is necessary to delineate the viral regulation of cGAS/STING signaling and their role in facilitating the KSHV lytic life cycle.

In this study, we focus on KSHV ORF48, a largely uncharacterized KSHV protein previously identified as a negative regulator of the cGAS/STING pathway, based on a luciferase screening assay in HEK293T cells(10). ORF48 was also found to interact with the PPP6 complex, which acts as a negative regulator of STING-dependent innate immune signaling(29). ORF48 is conserved among other gamma-herpesviruses, such as MHV68 and EBV. Previous studies have found that MHV68 ORF48 is essential for efficient viral replication in vitro and in vivo(30). Moreover, EBV ORF48 homolog BRRF2 was shown to be important for optimal infectious virion production(31,32). This raises the question as to the importance of ORF48 in maintaining an optimal KSHV lytic cycle, and whether the mechanism is cGAS-STING dependent.

We utilized both a siRNA knockdown approach and a genetic deletion system to study the role of ORF48 in the KSHV lytic cycle (33–35). Our results show that loss of ORF48 at either the mRNA level or gDNA level represses the mRNA and protein levels of multiple KSHV lytic genes and causes attenuated viral production. In addition, ORF48 removal at the gDNA level results in more intensive and global repression of the KSHV transcriptome, likely through disruption of RTA promoter activity. Collectively, these data highlight the importance of maintaining the integrity of ORF48 at both the gDNA and mRNA levels. At the protein level, we found that expressed ORF48 can interact with endogenous STING in HEK293 cells. Moreover, ORF48-stable HUVEC cells (HUVEC-ORF48) responded less to STING agonist treatment than EV-stable HUVEC cells, demonstrating the role of ORF48 in repressing STING function. Consistently, the removal of ORF48 in the HUVEC-ORF48 cell line resulted in elevated IFNβ production upon STING agonist stimulation. However, we did not observe a significant induction of IFNβ transcription in the absence of ORF48 during KSHV lytic reactivation, which is explained by the expression of other KSHV viral factors that have been shown to redundantly repress this pathway. Overall, our data suggest that the integrity of ORF48 is essential for optimal KSHV lytic replication, through multiple mechanisms.

## Materials and Methods

### Cell culture and reagents

iSLK.BAC16 (WT, delORF48#1, and delORF48#4), iSLK.219, iSLK.RTA, HEK293 and HEK293T cell lines were cultured in Dulbecco’s modified Eagle’s medium (DMEM), supplemented with 10% fetal bovine serum, and 1% penicillin-streptomycin). iSLK.BAC16 cells were cultured in the presence of 1 μg/ml puromycin, 250 μg/ml neomycin and 1.2 mg/ml hygromycin. iSLK.RTA cells were cultured in the presence of 1 μg/ml puromycin and 250 μg/ml neomycin. iSLK.219 cells harboring latent rKSHV.219 were maintained in DMEM supplemented with 10% FBS, 1% penicillin/streptomycin, G418 (250 ug/ml), hygromycin (400 ug/ml), puromycin (10 ug/ml). HUVEC-derived cell lines were cultured in EGM2 media from Lonza. All cells were maintained at 37 °C in a 5% (vol/vol) CO2 laboratory incubator subject to routine cleaning and decontamination. Interferon stimulatory DNA (ISD)(36) was synthesized from Eurofins company, ISD (sense), TACAGATCTACTAGTGATCTATGACTGATCTGTACATGATCTACA; ISD-reverse was the reverse sequence of above. An equal molar of ISD and its antisense oligos were annealed in PBS at 75°C for 30 min before cooling to room temperature overnight. STING agonist diABZI was purchased from MedchemExpress (HY-112921A). The plasmids pCDNA4.TO-ORF48-2xCSTREP, pCDNA4.TO-ORF39-2xCSTREP, and pCDNA4.TO-ORF37-2xCSTREP are kind gifts from Dr. Britt Glaunsinger (37), and can also be obtained from Addgene #136209, #136200 and #136198. The RTA expressing plasmid and RTA promoter plasmids were kindly provided by Dr. Zsolt Toth (38). The HA-STING plasmid is a kind gift from Dr. Glen Barber’s lab (39).

### Western blot and Immunoprecipitation

Antibodies used: mouse anti-viral interleukin-6 (vIL6) antibody(40) is a kind gift from Dr. Blossom Damania. The anti-KSHV ORF26 (MA5-15742), anti-KSHV ORF45 (MA5-14769) and anti-STREP-TAG II (MA5-37747) were obtained from Invitrogen. The anti-KSHV ORF57-HRP (sc-135746), anti-KSHV K8.1 A/B-HRP (SC-65446) and anti-human actin-HRP (sc-47778) were purchased from Santa Cruz. The anti-TBK1 (38066S), anti-phospho-TBK1 (5483S), anti-IRF3 (11904S), anti-HA-tag (3724s), anti-FLAG-HRP (86861s) and anti-STING (13647S) were purchased from Cell Signaling. The anti-phospho-IRF3 (AB76493) antibody was obtained from Abcam. The rabbit anti-ORF48 polyclonal antibody was generated from the Abclonal company.

### siRNA transfections and KSHV reactivation analyses in iSLK.219 cells

iSLK.219 cells were maintained as previously described and were transfected with TransIT-X2 (Mirus MIR6004) according to the manufacturer’s specifications. At 24 hours post-transfection, cells were treated with doxycycline (Dox, 0.2 ug/ml) for KSHV lytic reactivation. Cells and supernatant were collected at 0h, 24h, 48h, and 72 hours post-reactivation.

siRNAs were synthesized by Sigma with the following designed sequences:

siNS: *UGGUUUACAUGUCGACUAA*

si*ORF48*#5: *GGUGAUGCAAUUAGAGAAA*

si*ORF48*#6: *UGGGAUGACUGCAAAGAUA*

### KSHV genome array

We utilized a modified system as previously described by other groups, with newly designed primers (10,27). Briefly, two to four sets of RT-PCR primers based on the sequence of each KSHV ORF were re-designed and the most specific primer set with the lowest background in untreated iSLK.219 and highest fold induction in Dox-treated iSLK.219 was selected for each ORF. RNAs from each group were extracted from duplicate samples to synthesize cDNA. Eighty-eight KSHV viral transcript levels were analyzed using a real-time qPCR-based KSHV transcriptome array. mRNA levels of viral genes were normalized to the mRNA levels of *GAPDH* to yield dCT as a measure of relative expression. These were then subjected to unsupervised clustering. A heat map and dendrogram depicted by the brackets is shown. Higher transcript expression levels are indicated by red and lower expression levels by blue as shown in the key.

### KSHV constructs and establishment of stable iSLK.BAC16 cells harboring ORF48 deletion

The detailed protocol is as previously described(35). Briefly, primers were designed to amplify the Kanamycin (Kan)-resistance gene with homology to KSHV sequences upstream and downstream of ORF48. BAC16-ORF48del forward primer: *GGAAGACGATGGGGGAAATGTGGCATT- ACCTGACACGGTTGTTCAGTCACATGTACGCTA-AGGATGACGACGATAAGTAGGG*, reverse primer: *GGGGTTGGGTGGGGAGACCCTAGCGTACATGTGACTGAACAACCGTGTCAGGTA- ATGCCAAACCAATTAACCAATTCTGATTAG*. Upon electroporation, the Kan-cassette is inserted into BAC16 and a Kan-resistant bacmid is generated. Treatment with the I-SceI enzyme results in the linearization of the bacmid, allowing intramolecular recombination, which generates the final bacmid without Kanamycin and with the deletion of ORF48. All BACmid mutants and one WT BAC16 BAC mid were digested with NheI and subject to restriction fragment length polymorphism (RFLP) analysis based on the PFGE system. Two clones ORF48del#1 and #4 were selected and validated with sequencing. The genetically modified BAC16 (ORF48del#1 and ORF48del#4) were transfected into HEK293T cells, which were selected with hygromycin for approximately 2 weeks, and treated with sodium butyrate (NaBr) and 12-O-tetradecanoylphorbol-13-acetate (TPA) to induce virus production. Then, at 72 hours post-induction, the supernatants containing viruses were collected, filtered, and utilized to infect iSLK.RTA cells. Positive iSLK.BAC16 cells were selected with puromycin, hygromycin, and neomycin. BAC16+ iSLK cells were also visually tracked using green fluorescent protein (GFP) expression since BAC16 contains a GFP cassette under the regulation of the constitutive promoter EF-1a.

### Viral genome copy quantification and viral infection assay

KSHV genome copies were quantified as previously described(27). Briefly, gDNA from cells or supernatants were purified with the DNeasy blood and tissue kit (Qiagen). The pCDNA4.TO- ORF39-2xCSTREP plasmid(37) was utilized to generate a standard curve for the cell cycle threshold (CT) versus the genome copy number. The primers used to amplify the genome of KSHV were located in the ORF39 region. ORF39 F: 5’-*GTGGGAGTATTCGTGGGTTATC*-3’; R: 5’-*GGTGAACAGTCGGAGTTCTATC*-3’. Supernatants collected from reactivated iSLKs (iSLK.BAC16 or iSLK.219) were utilized to infect naïve HEK293T cells, supplemented with 8 ug/ml Polybrene. Spin-inoculation was performed by centrifuging the plates at 2500 RPM, 30°C, for 90 minutes. Genomic DNA from the infected cells was extracted for Viral genome copy quantification.

### RT-PCR

Total RNA was isolated with the RNeasy extraction kit (Qiagen), and cDNA was synthesized with a cDNA synthesis kit (MedchemExpress #HY-K0510), according to the manufacturer’s protocol. qPCR was performed using the SYBR Green qPCR master mix (MedchemExpress #HY-K0501), previously mixed with ROX reference dye II. Primers used for SYBR green qRT-PCR were:

KSHV gene primers:

*ORF57* F: 5’-*TGGACATTATGAAGGGCATCCTA*-3’; R: 5’-*CGGGTTCGGACAATTGCT*-3’.

*ORF39* F: 5’- *GTGGGAGTATTCGTGGGTTATC*-3’; R: 5’-*GGTGAACAGTCGGAGTTCTATC*-3’.

*K8.1* F: 5’-*AAAGCGTCCAGGCCACCACAGA*-3’; R: 5’-*GGCAGAAAATGGCACACGGTTAC*-3’.

*ORF48* F: 5’-*TGATCTGGGATGACTGCAAAG-*3’; R: 5’-*AAAGAATGTGTCTCCCGTGG*-3’.

Human gene primers:

*GAPDH* F: 5’-*GTCTCCTCTGACTTCAACAGCG*-3’; R: 5’-*ACCACCCTGTTGCTGTAGCCAA*-3’.

*IFNβ* F: 5’-*AGTAGGGCGACACTGTTCGTG*-3’; R: 5’-*GAAGCACAACAGGAGAGCAA*-3’.

The relative amount of IFNβ, ORF48, ORF57, ORF36, and K8.1 mRNA was normalized to *GAPDH* RNA level in each sample, and the fold difference between the treated and mock samples was calculated.

### Luciferase assay

The plasmids were obtained from Dr. Zsolt Toth and the detailed protocol was followed as previously described (38). Briefly, HEK293T cells were co-transfected with RTA-promoter luciferase plasmids, an RTA expression plasmid, and a CMV-Renilla plasmid using Mirus Transit X2 (Mirus MIR6004). At 48 hours post-transfection, the luciferase assay was performed using a luciferase assay kit from Promega following the manufacturer’s instructions. Each luciferase experiment was performed at least three times, and three biological samples per treatment were used. Results were generated as a ratio of Firefly/Renilla luminescent intensity.

### Statistical analysis

Statistical significance of differences in cytokine levels, mRNA levels, viral titers, and luciferase intensity in reporter assay were determined using Student’s t-test. * indicates P<0.05, ** indicates P<0.01, *** indicates P<0.001, **** indicates P<0.0001.

## Results

### Knockdown of ORF48 attenuates KSHV lytic replication

To study the role of ORF48 during the KSHV lytic cycle, we utilized siRNA to knock down *ORF48* mRNA in iSLK.219 cells. As shown in Figure 1A, iSLK.219 cells were transfected with *ORF48* specific siRNAs (or non-scramble control siRNA, NS) for 24 hours, followed by treatment with doxycycline for 24h, 48h, and 72h. The iSLK.219 system features a dual indicator system, in which GFP is expressed as an indicator of latency and RFP reflects lytic reactivation status (34). As shown in Figure 1B, fewer positive cells and less RFP intensity were observed in the si*ORF48* group than in the siNS group. The fluorescence intensity of each well was scanned and quantified by a plate reader to generate the RFP/GFP ratio under each condition. Consistently, we observed a reduced RFP/GFP intensity ratio upon *ORF48* knockdown, indicating a reduced lytic reactivation status in these cells (Figure 1C). Next, we assessed the impact of ORF48 knockdown in the expression of KSHV lytic genes. We chose three KSHV genes *ORF57*, *ORF39*, and *K8.1*, as representative genes transcribed at immediate early (IE), early (E), and late (L) stages respectively (43). Knockdown of *ORF48* failed to repress *ORF57* (IE) transcription (Figure 1D), while the *ORF48* knockdown group showed less *ORF39 (E)* transcription at 24 hours (Figure 1E) and significantly reduced *K8.1* (L) transcription at 72 hours (Figure 1F). In addition, we found that with less ORF48, KSHV failed to replicate its genome as efficiently as in siNS-treated groups (Figure 1G). We also measured the protein expression levels of multiple KSHV ORFs from different lytic stages. ORF48 expression was detected as early as 24 hours, and was successfully knocked down at all time points, consistent with the previous report defining ORF48 as an immediate early gene (41,42). ORF57 from the IE stage was not downregulated with less ORF48 expression. Less ORF45 and vIL6 were detected upon ORF48 knockdown at 24 hours, but quickly recovered to a similar level to the siNS-treated group. The late-stage expressed ORF26 and K8.1 were repressed after *ORF48* knockdown, especially at 72 hours after infection (Figure 1H). These protein expression levels are consistent with the transcript levels for each gene as shown in our RT-PCR data. Additionally, significantly fewer KSHV genome copies were detected in the supernatant of si*ORF48*-treated iSLK.219 cells (Figure 1I). Infection assay confirmed that the si*ORF48* group produced fewer infectious virions (Figure 1J and K). In general, we observed that late gene expression and virion release rely upon optimal ORF48 expression.

**Figure 1.**
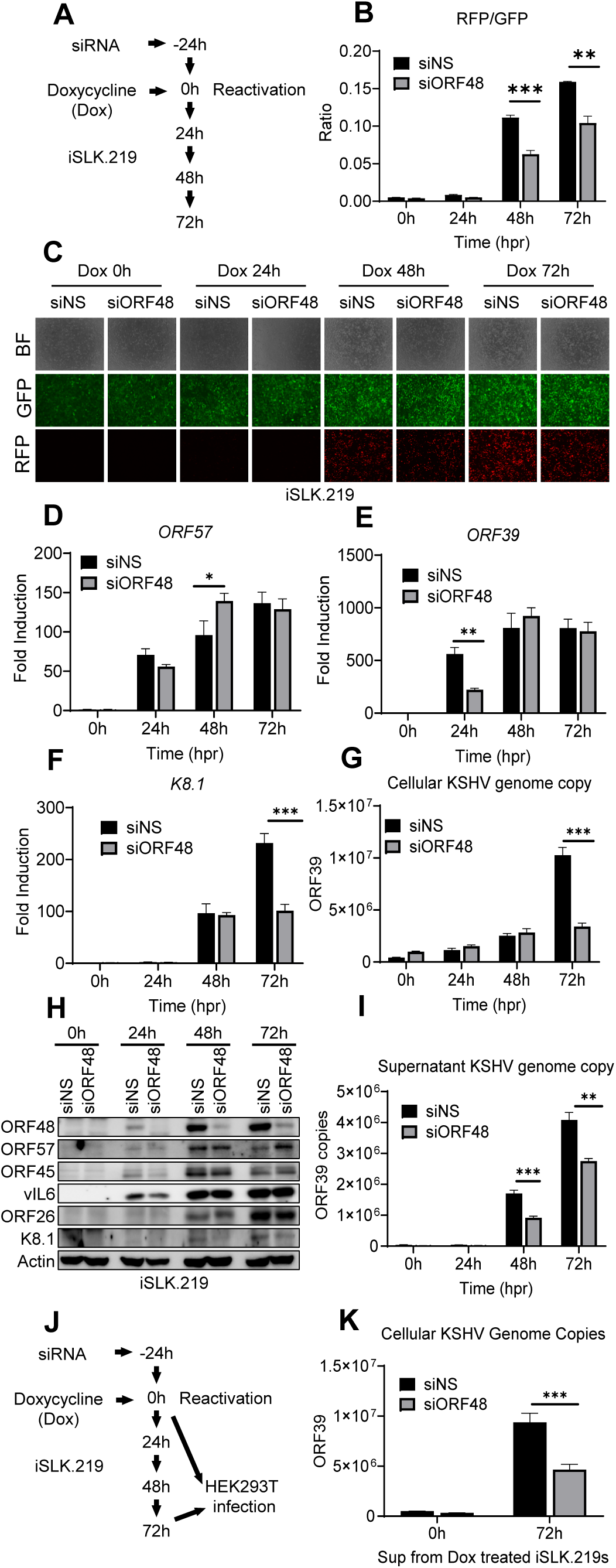
Knockdown of *ORF48* attenuates KSHV lytic replication. (A) Schematic illustration of the experimental procedure of (B-H). (B) The RFP and GFP fluorescence intensity were measured in groups at each time point. Briefly, a scan mold of the plate reader will read 21 spots spreading in each well to calculate the average fluorescence intensity. (C) Representative microscope image of bright field, GFP, and RFP in each group. (D-F) Total RNAs were extracted from all groups at all time points to synthesize cDNA, and subjected to RT-PCR. Specific RT-PCR primers were used to detect (D) *ORF57* representing an immediate early lytic gene, (E) *ORF39* representing an early lytic gene, and (F) *K8.1* representing a late lytic gene. Expression levels of these genes were normalized with GAPDH. (G) Cellular KSHV genome copies were quantitated using a genomic primer based on the ORF39 coding sequence as previously described. A STREP-tagged ORF39 (37) was used to generate the standard curve. (H) Western blot analysis of ORF57, vIL6, and ORF45 (immediate early or early stage); ORF26 and K8.1 (late stage). (I) The supernatants from all groups containing KSHV genome copies were quantitated using the same method as (G). (J) Schematic illustration of the experimental procedure of infection assay. Briefly, the supernatants from 72h groups were collected to infect naïve HEK293T cells to evaluate infectious virion productions from each group. Zero-hour groups served as a negative control for infection. (K) Forty-eight hours post-infection, genome DNAs from infected HEK293 cells were extracted and the KSHV genome copy numbers were evaluated by the same method as (G). Data are presented as mean ± s.d. from at least three independent experiments. *indicates p<0.05. ** indicates p<0.01 *** indicates p<0.001 **** indicates p<0.0001 by Student’s t-test.

### Knockdown of ORF48 inhibits KSHV lytic gene transcription

Since inhibitory patterns on selected ORF mRNA levels were observed previously, we further evaluated this pattern on the KSHV transcriptome. We utilized a modified system as previously described by other groups (10,27), but with newly designed primers. We measured the KSHV transcriptome as described in Figure 1A, by harvesting RNA, and subjecting the cDNAs of all groups to the RT-PCR-based KSHV transcriptome array at each time point upon reactivation. Generally, the si*ORF48* groups exhibited attenuated and delayed transcription on most KSHV ORFs, and this impact was most obvious at 72 hours (Figure 2A). We then calculated the mean expression level of each gene at every condition and generated normalized ratios in the form of (si*ORF48* expression)/(siNS expression). Upon ORF48 knockdown, the majority of KSHV genes were distributed around a ratio of one at 24h, but aggregated to a mean ratio of approximately 0.8 at 72h (Figure 2B). We further categorized KSHV genes into four sets, upregulated (ratio>1.2), not changed (0.8-1.2), downregulated (0.4-0.8) and highly downregulated (<0.4). At 24h and 48h, a very small portion of KSHV ORFs were upregulated, about half of the genes were not affected (0.8-1.2) and slightly less than half of the genes were downregulated (0.4-0.8). However, at 72h, the majority of the genes were downregulated and only a few of the genes remained unaffected (Figure 2C). This is consistent with our findings that loss of ORF48 seems to have a more profound impact at a later stage of the KSHV lytic cycle.

**Figure 2.**
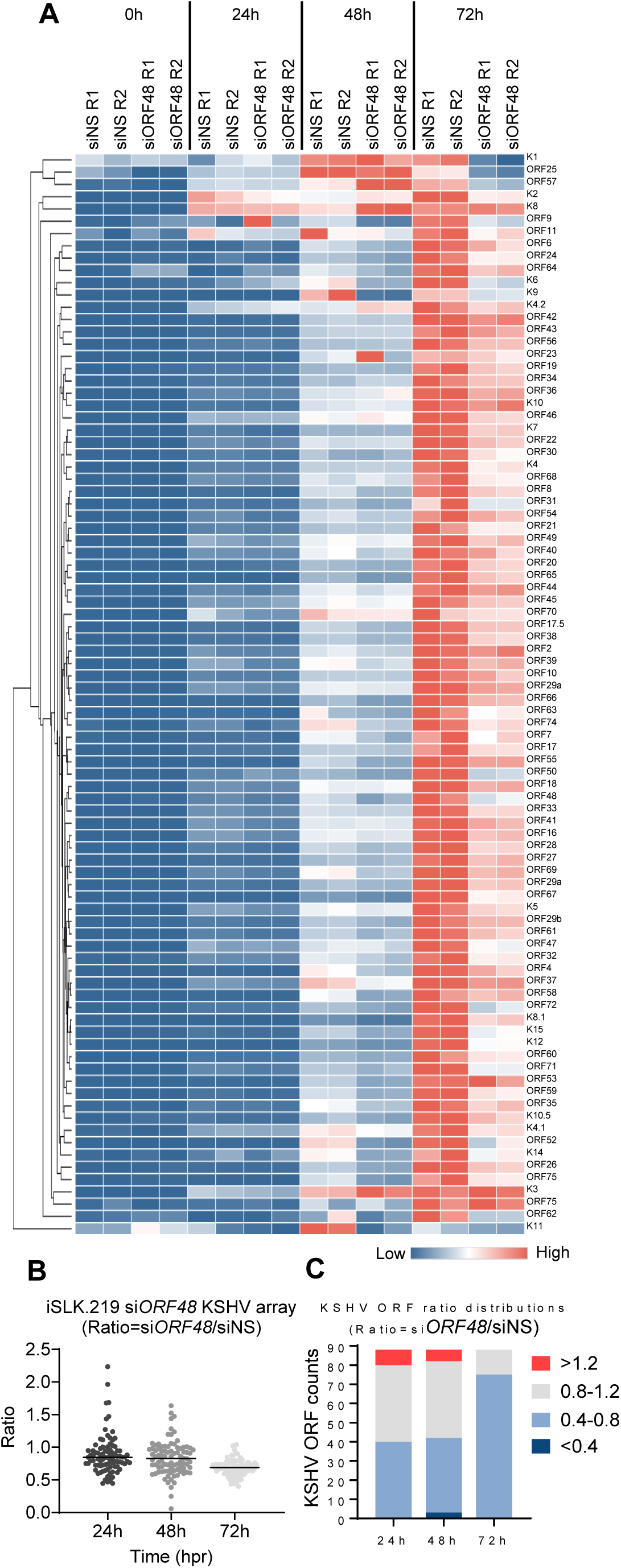
Knockdown of ORF48 inhibits KSHV lytic gene transcription. (A) iSLK.219 transfected with siNS or siORF48 treated as described in the text and Figure 1A. A real-time qPCR-based KSHV transcriptome array was performed. Higher transcript expression levels are indicated by red and lower expression levels by blue as shown in the key. (B) For each KSHV ORF at 24h, 48h, and 72h time points, the average expression level of two biological replicates of si*ORF48* was normalized to their siNS controls to generate a ratio. The plot depicts summary statistics and the density of each KSHV ORF from each group, and each dot represents one of the eighty-eight KSHV ORFs. (C) The distribution of KSHV ORF ratios in each group was further categorized into upregulated (>1.2), unaffected (0.8-1.2), downregulated (0.4-0.8) and highly downregulated (<0.4).

### Construction of BAC16-ORF48 deletion mutants using the BAC16 system

The integrity of genomic DNA is the foundation of appropriate mRNA and protein expression. Thus, we decided to genetically remove ORF48 from KSHV and build an iSLK cell line carrying this ORF48 deletion mutant or WT KSHV. This would allow us to 1) study the role of ORF48 in the KSHV lytic life cycle in more stringent conditions and 2) evaluate the importance of ORF48 genomic integrity. We utilized the KSHV BAC16 system, which carries the complete KSHV genome and enables its genetic manipulation in E. coli. As previously described (35), we used the Bacterial Artificial Chromosome-based two-step bacteriophage lambda Red-mediated recombination system. Since there is no overlapping gene encoding region to adjacent genes, ORF47, ORF49 and RTA, we decided to remove the entire ORF48 coding sequence to ensure the complete abolishment of ORF48 expression (Figure 3B). BACmids were digested with NheI and subject to restriction fragment length polymorphism (RFLP) analysis based on the PFGE system. As shown, the BAC16-ORF48-del mutants maintain the integrity of the KSHV genome, except for one predicted band shift from 25,693bp to 24,484bp due to the loss of ORF48 coding sequence (Figure 3A, 3C). We picked two ORF48 deletion mutant clones, ORF48del#1 and ORF48del#4, and their sequences were verified by DNA sequencing. As previously described, we then used the WT as well as ORF48 deletion BACmids to generate iSLK.BAC16, to study the role of ORF48 in the KSHV lytic life cycle. To attenuate the influence of genetic instability during iSLK.BAC16 stable cell generation, we created two stable cell lines iSLK.BAC16-ORF48del#1 and #4. A fresh iSLK.BAC16 WT cell line was also generated simultaneously to serve as a control for the following experiments (Figure 3D).

**Figure 3.**
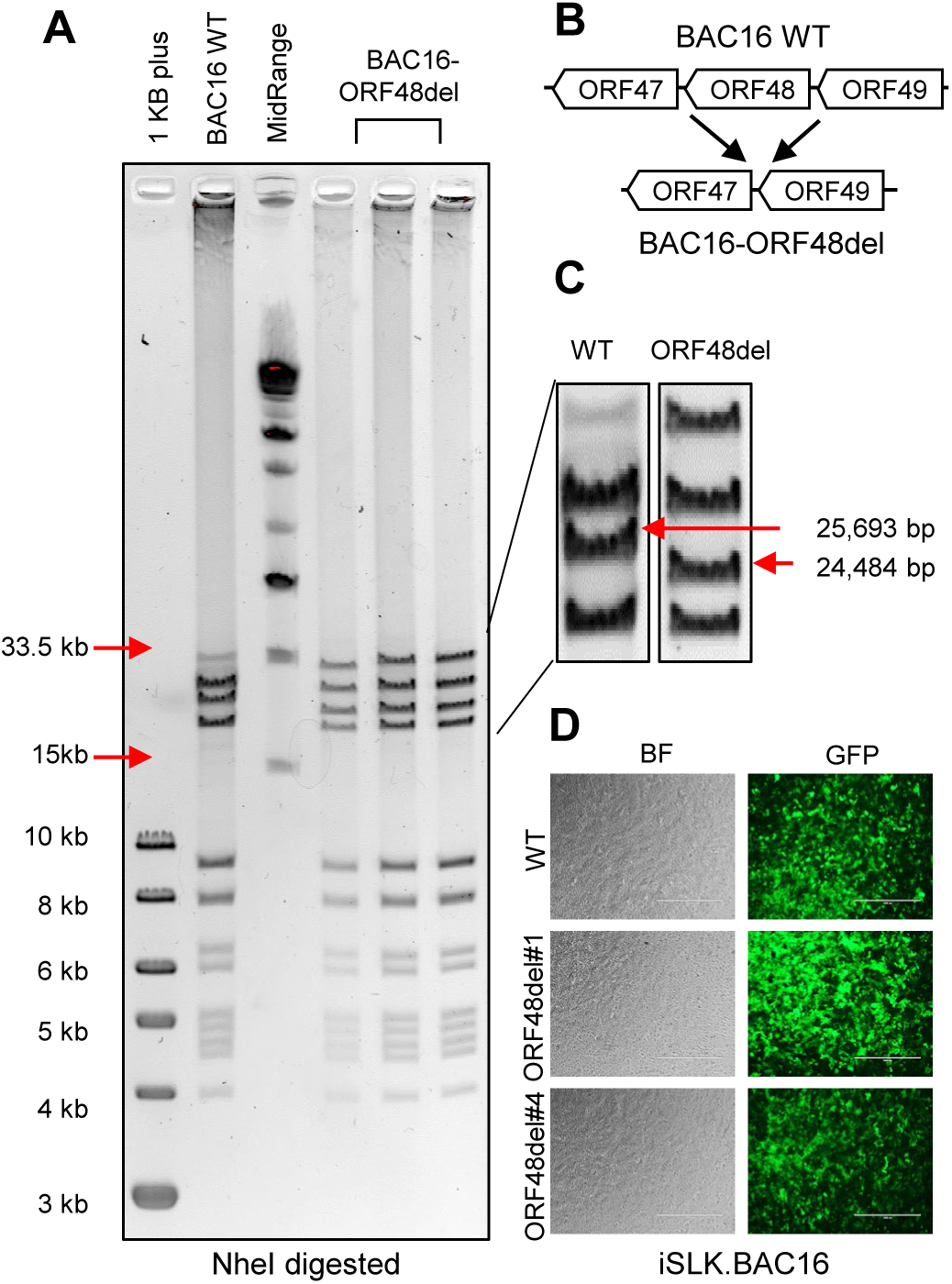
Construction of ORF48 deletion mutants using the BAC16 system. (A) Agarose gel electrophoresis of wild-type BAC16 and mutant BAC16-ORF48del. The loading sequence from left to right is 1kb plus DNA ladder, WT BAC16, midrange DNA ladder, BAC16-ORF48del candidates #1, #4 and #6. DNA was digested with NheI for two hours and resolved on a 0.4% agarose gel stained with ethidium bromide. DNA ladder sizes covering the KSHV genome are indicated to the left of the gels. (B) Schematic illustration of the strategy used to generate ORF48 deletion mutants. The ORF48 coding region does not overlap with adjacent ORF47 and ORF49. (C) Analysis of BAC16-ORF48del VS WT BAC16. Deletion of the ORF48 coding region (1.2kb) results in a decreased size of a band from 25,693bp to 24,484bp, as indicated by red arrows. (D) Bright-field view and GFP expression in the established WT, ORF48del#1 and #4.

### ORF48 deletion significantly defects KSHV lytic replication

Upon establishing three iSLK cell lines carrying either iSLK.BAC16 WT, ORF48del#1, or ORF48del#4, we next aimed to explore the role of ORF48 on the KSHV lytic cycle. We treated these cell lines with doxycycline, which induces ORF50 (RTA) expression in cells, a necessary and sufficient event to trigger KSHV lytic reactivation. At 0h, 24h, 48h, and 72h after reactivation, lytic replication statuses were evaluated in these three groups (Figure 4A). We first confirmed that *ORF48* transcriptions were completely abolished in both del#1 and del#4 mutants (Figure 4B). ORF48 deletion significantly suppressed the mRNA level of ORF57 (IE), ORF39 (E) and K8.1 (L) genes (Figure 4C-E). Moreover, ORF48 deletion mutants showed attenuated KSHV genome replication in comparison with WT BAC16 upon reactivation (Figure 3F). Consistently, ORF48 deletion groups expressed ORF57, vIL6, and ORF45 significantly less than the WT group (Figure 3G). We further quantified the KSHV genome copy number representing virion production in these three groups. The removal of ORF48 resulted in significantly fewer KSHV genome copies in both ORF48 deletion groups (Figure 3H). We used the same volume of the supernatants to infect naïve HEK293T cells (Figure 3I). Consistently, we detected less KSHV genome in HEK293T cells infected with ORF48 deletion group generated supernatants, indicating less infectious virions were produced from these iSLKs harboring ORF48 deletion mutants (Figure 3J). In all, these data suggested a fundamental role of ORF48 in the KSHV lytic cycle, and its requirement for optimal virion production and viral propagation.

**Figure 4.**
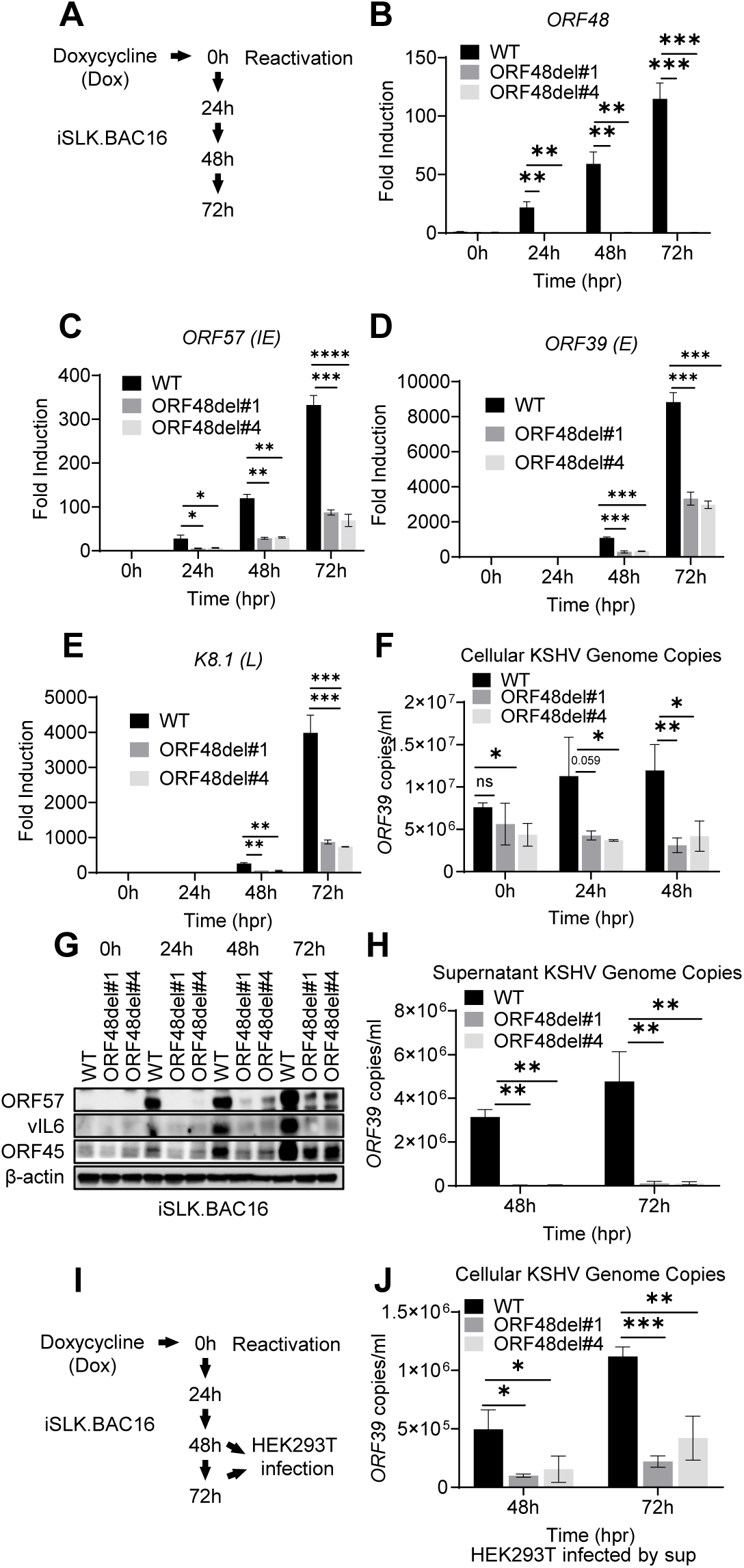
ORF48 deletion defects KSHV lytic replication. (A) Schematic illustration of the experimental procedure of (B-H). Total RNAs were extracted from all groups at all time points to synthesize cDNA and subjected to RT-PCR. Specific RT-PCR primers were used to detect (B) ORF48 for deletion confirmation, (C) *ORF57* representing an immediate early lytic gene, (D) *ORF39* representing an early lytic gene, and (E) *K8.1* representing a late lytic gene. Expression levels of these genes were normalized with *GAPDH*. (F) Cellular KSHV genome copies were quantitated using a genomic primer based on the ORF39 coding sequence as previously described. A STREP-tagged ORF39 (37) was used to generate the standard curve. (G) Western blot analysis of ORF57, vIL6 and ORF45 encoded by KSHV. (H) The supernatants from all 48h and 72h groups containing KSHV genome copies were quantitated using the same method as (F). (I) Schematic illustration of the experimental procedure for infection assay. Briefly, the supernatants from all 48h and 72h groups were collected to infect naïve HEK293 cells to evaluate infectious virion production. (J) Forty-eight hours post-infection, genome DNAs from infected HEK293 cells were extracted and the KSHV genome copy numbers were evaluated by the same method as (F). Data are presented as mean ± s.d. from at least three independent experiments. *indicates p<0.05. ** indicates p<0.01 *** indicates p<0.001 **** indicates p<0.0001 by Student’s t-test.

### ORF48 genome deletion causes global KSHV ORF transcriptional repression

Since ORF48 deletion caused inhibition on all representative viral genes or proteins, we further evaluated if this impact has some specificity or is global for the entire KSHV transcriptome. The same experiments were performed as shown in Figure 4A, and the samples were subjected to the RT-PCR-based KSHV transcriptome array at each time point upon reactivation. The depletion of ORF48 led to a massive suppression of nearly all KSHV gene transcriptions at each time point (Figure 5A). We calculated the mean expression level of each gene at every condition and generated normalized ratios in the form of (ORF48del#1 or #4 expression)/(WT expression). As seen in Figure 5B, upon ORF48 deletion, the mean ratios of all groups were less than 0.4 (Figure 5B). We then used the same categorization standard as described in Figure 2C. In all groups in each time point, the majority of the genes fall into the category of highly downregulated, a decent number of genes are downregulated, while only a few ORFs remain unaffected or upregulated (Figure 5C). While we expected a stronger phenotype using the genetic deletion model, the data that ORF48 deletion caused such an early and robust disruption of the KSHV lytic cycle still drew our attention. Particularly, we are curious about the additional potential impact caused by comprised ORF48 genome integrity.

**Figure 5.**
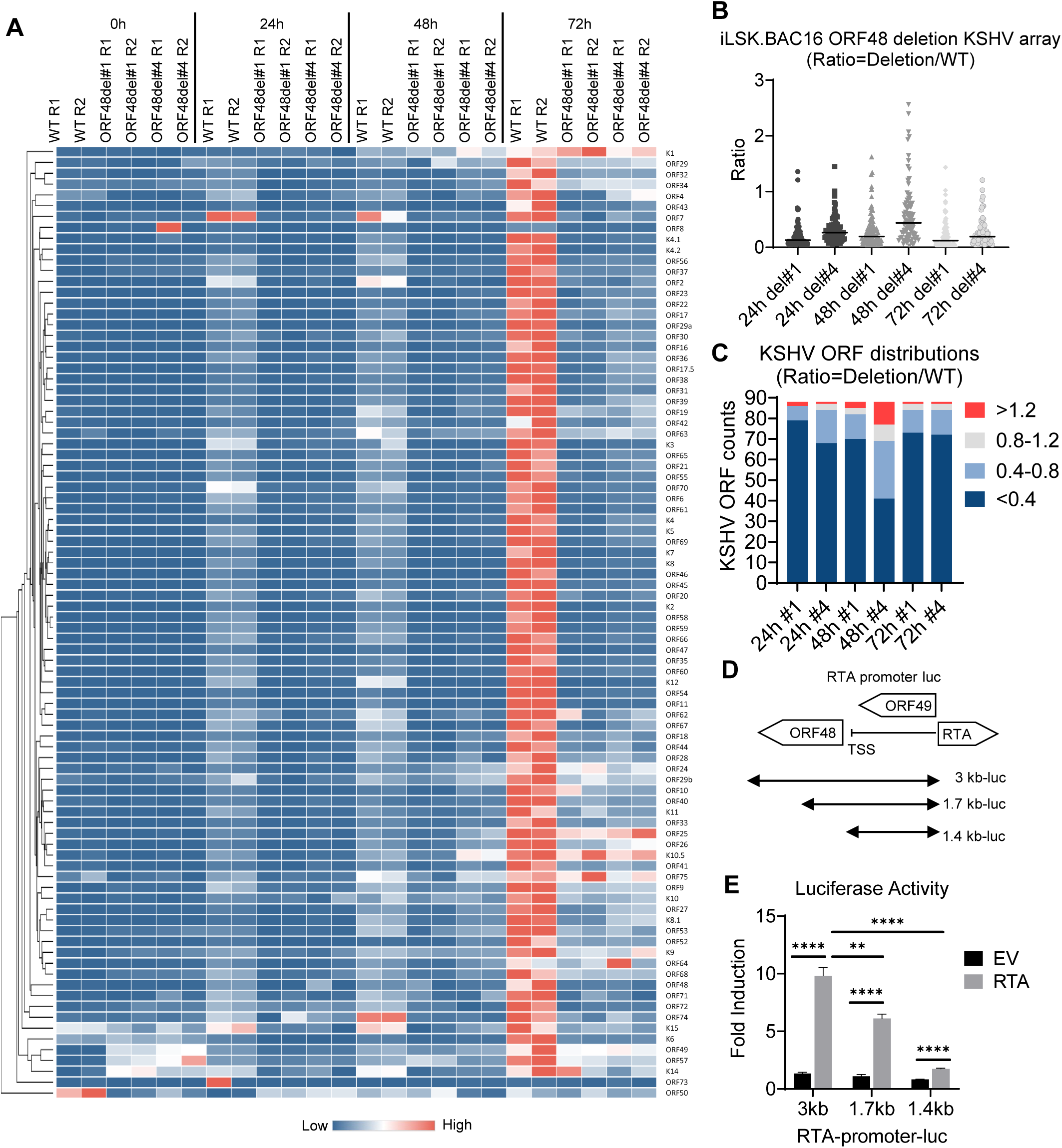
ORF48 genome deletion causes global KSHV ORF transcriptional repression. (A) iSLK.BAC16 WT, ORF48del#1 and ORF48del#4 cells were treated as described in the text and Figure 3A, and subject to the KSHV transcriptome array as described in Figure 2A. Higher transcript expression levels are indicated by red and lower expression levels by blue as shown in the key. (B) The average expression level of two biological replicates of ORF48del#1 or ORF48del#4 was normalized to their WT controls to generate a ratio. The plot depicts summary statistics and the density of each KSHV ORF from each group, and each dot represents one of the eighty-eight KSHV ORFs. (C) The distribution of KSHV ORF ratios in each group was further categorized into upregulated (>1.2), unaffected (0.8-1.2), downregulated (0.4-0.8) and highly downregulated (<0.4). (D) Schematic diagram of RTA-promoter constructs. The 3 kb, 1.7 kb, and 1.4 kb upstream of the RTA coding sequence were cloned into the upstream of the firefly luciferase reporter. (E) The RTA-promoter constructs and a CMV-renilla luciferase construct were co-transfected with pCDNA-FLAG-RTA plasmid or empty vector control into HEK293T cells. Forty-eight hours later, cells were harvested, lysed, and subjected to a Dual-luciferase assay. Firefly/Renilla ratios were generated in each group and all groups were then normalized to their EV control group respectively to generate fold induction. Data are presented as mean ± s.d. from at least three independent experiments. *indicates p<0.05. ** indicates p<0.01 *** indicates p<0.001 **** indicates p<0.0001 by Student’s t-test.

The KSHV genome shows no overlapping of ORF48 with adjacent ORFs, and the ORF49 start codon and RTA start codon are both approximately 1.3 Kb away from the ORF48 start codon. However, we did notice that the ORF48 coding region is only slightly over 200 bp away from the transcription start site (TSS) of RTA (ORF50), which transcribes in the opposite direction of the KSHV genome. Given the proximity of ORF48 to the RTA TSS, the removal of the ORF48 genome sequence may affect RTA promoter activity, which plays a pivotal role in KSHV lytic reactivation(6,43). To test this hypothesis, we evaluated the impact of different RTA promoter lengths on RTA-dependent self-promoter activation. As shown in Figure 5I, we obtained three RTA promoter-luciferase constructs covering the whole (3 kb from RTA start codon), partial (1.7 kb from RTA start codon), or none (1.4 kb from RTA start codon) ORF48 genome region. As seen in Figure 4D, the transfected RTA plasmid successfully activated the RTA promoter in the 3 kb group compared with the EV-transfected group. A loss of RTA promoter activity was observed when the ORF48 coding region was partially removed from the RTA promoter. Especially in the 1.4 kb group, which mimics the ORF48 deletion mutant conditions, RTA promoter activation was nearly at the basal level (Figure 5E). These data suggest that our previous observation in deletion mutants is due to both lack of ORF48 expression and compromised RTA-promoter activity. Collectively, loss of *ORF48* mRNA caused selective attenuation of KSHV gene transcription, while deletion of ORF48 encoding DNA led to global KSHV transcriptomic repression. All of these indicate the crucial role of maintaining the integrity of ORF48-encoding DNA and the expression of ORF48 in an optimal KSHV lytic cycle.

### ORF48 interacts with STING and blocks STING-dependent innate immunity

We next probed for the mechanism by which ORF48 expression is important for KSHV lytic reactivation from latency. Previously, ORF48 was identified as one of the negative factors in our cGAS-STING reconstitution-based IFNβ promoter assay. Therefore, we first explored if ORF48 interacts with STING. As shown in Figure 6A, overexpressed STREP-ORF48 was co- immunoprecipitated with overexpressed HA-STING, while overexpressed STREP-ORF37 (control) failed to bind STING in HEK293T cells. We also detected endogenous STING interacting with STREP-ORF48 in HEK293 cells (Figure 6B), building a potential connection between ORF48 and STING function. To better evaluate this, we established a FLAG-tagged ORF48 stable cell line in telomerase-immortalized HUVEC. An empty vector stable cell line was also created simultaneously as a negative control. We treated these two cell lines with ISD to stimulate cGAS and analyzed the differences in innate immune response. As shown in Figure 6C, HUVEC-ORF48 failed to mount a similar level of IFNβ as HUVEC-EV. Consistently, less p-TBK1 and p-IRF3 were observed in HUVEC-ORF48 at both 3 hours and 6 hours after ISD transfection. Expressions of ORF48 were detected only in HUVEC-ORF48 cells (Figure 6D). We observed a similar pattern in the STING agonist diABZI-treated experiments (Figure 6E-F), suggesting that ORF48 alone blocks STING signaling in our stable cell-based system. Therefore, we hypothesized that ORF48 is required for optimal KSHV lytic cycle through blocking STING-dependent innate immune signaling. However, when we evaluated ORF48’s role in the KSHV reactivation system, we failed to observe significant IFNβ transcription induction in cells expressing less ORF48 (Figure 6G). In addition, we did not observe upregulations of p-TBK1 or p-IRF3 in si*ORF48*-treated cells, compared with the siNS group (Figure 6H). We next tested the loss of function of ORF48 in our virus-free HUVEC-ORF48 stable cell line system. Indeed, the knockdown of ORF48 enhanced IFNβ production and p-TBK1/IRF3 upon diABZI treatment (Figure 6I-J). These data suggest that during the KSHV lytic cycle, the loss of ORF48 is compensated by other viral negative regulators of STING signaling, and therefore no significant increase of IFNβ was observed. Although standalone ORF48 is capable of blocking STING-dependent signaling, this function is redundant in our iSLK-based system. Therefore, in the iSLK system, ORF48 is required for optimal KSHV lytic replication through an unknown mechanism that is independent of STING-based innate immunity.

**Figure 6.**
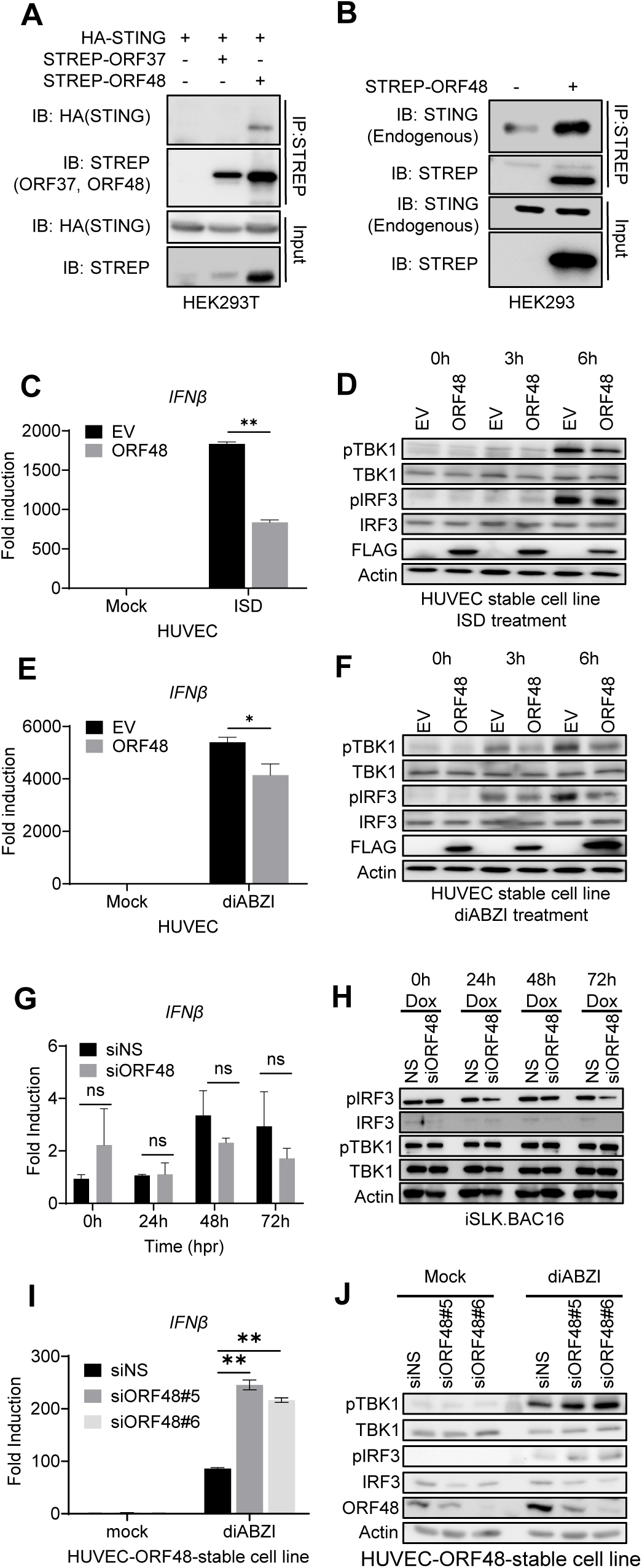
ORF48 interacts with STING and blocks STING-dependent innate immunity. (A) Co-immunoprecipitation of HA-STING and STREP-ORF48. HEK293T cells were transfected with HA-STING, an empty backbone, STREP-ORF37, or STREP48 as shown. Forty-eight hours later, cell lysates were immunoprecipitated with STREP antibody and protein A/G beads. HA or STREP antibodies were used for band detection. (B) Co-immunoprecipitation of endogenous STING and STREP-ORF48. HEK293 cells were transfected with an empty backbone or STREP-ORF48 as shown. Forty-eight hours later, cell lysates were immunoprecipitated with STREP antibody and protein A/G beads. STING or STREP antibodies were used for band detection. (C-F) HUVEC-EV or HUVEC-ORF48 stable cells were transfected with ISD or diABZI for 0, 3, and 6 hours. RT-PCR of IFNβ in each group at six hours was performed as shown in (C) ISD or (E) diABZI. Western blot assays of each group at three and six hours were shown in (D) ISD or (F) diABZI. (G) iSLK.219 cells were treated as described in Figure 5A, and IFNβ levels were detected using RT-PCR. (H) Western blot assays evaluating p-TBK1, TBK1, p-IRF3, and IRF3 levels in the above samples. Beta-actin serves as a loading control. (I-J) HUVEC-ORF48 stable cell lines were transfected with two siRNAs targeting ORF48 for forty-eight hours. Samples were then subjected to 4uM of diABZI for 6 hours, and detected with either (I) RT-PCR for IFNβ production or (J) Western blot assays for p-TBK1, TBK1, p-IRF3, IRF3 and ORF48 evaluation. Data are presented as mean ± s.d. from at least three independent experiments. *indicates p<0.05. ** indicates p<0.01 *** indicates p<0.001 **** indicates p<0.0001 by Student’s t-test.

## Discussion

The cGAS-STING pathway is a critical component of immunity in mammalian cells, which detects cytosolic DNA and induces a potent anti-viral response(44). Therefore, it is extensively targeted and repressed by multiple human pathogens, including KSHV(19). Identifying the viral inhibitors and understanding their inhibitory mechanisms of STING are important for revealing the mechanisms of immune evasion by KSHV. This study is a continuing effort to further characterize a predicted KSHV-encoded negative regulator of the cGAS/STING pathway, and study how it affects KSHV lytic replication, a critical step for promoting persistent KSHV infection in the host.

Proper expression of a protein requires integrity of both genome DNA and mRNA. To better evaluate both aspects of ORF48 on the KSHV lytic life cycle, we utilized two systems. A siRNA-based knockdown system to evaluate the role of intact ORF48 mRNA, and a BAC16-based genetic deletion system to further study the impact of a comprised ORF48 gDNA. In both systems, we observed impaired KSHV DNA replication and attenuated KSHV viral production, highlighting the role of ORF48 in optimizing KSHV lytic replication. Interestingly, while removing *ORF48* mRNA caused a selective pattern of KSHV transcriptome, ORF48 gDNA removal led to an intensive and global inhibition.

Further dissection of the RT-PCR-based whole KSHV ORF transcriptome analysis suggested that a number of the immediate early and early transcripts tend to be less affected by *ORF48* knockdown, while late transcripts are prone to be impaired upon loss of *ORF48*. This is consistent with the immunoblot assays showing that the expression levels of some immediate early and early genes, such as ORF57, ORF45, and vIL6, are slightly reduced in si*ORF48* groups. Conversely, the representative late genes, such as K8.1 and ORF26, are reduced in si*ORF48* groups, especially during 48 and 72 hours. This data suggests that ORF48 could play a role in early gene expression, which may create a negative impact on KSHV DNA replication. It is not surprising that the accumulation of these negative effects upon losing ORF48 expression leads to less infectious virions.

The transcript containing *ORF48* (KIE4.1) was first detected in the immediate early stage of the KSHV lytic cycle (41,42), this is consistent with our observation that ORF48 mRNA and protein were detected at 24h upon reactivation. These pieces of evidence support that ORF48 plays a critical role in the initiation of lytic KSHV replication and expression of a broad range of KSHV lytic transcripts. Upon further investigation of ORF48 deletion, we found that the ORF48 coding region does not overlap with any other known KSHV transcripts, but is required for optimal RTA promoter activity, a critical step for optimal lytic reactivation. In addition, a previously reported CHIP-on-chip analysis showed a high enrichment of the activating histone modifications in a region encoding IE protein ORF45, ORF48, and ORF50 (RTA). These findings suggested that this genomic region is critical for appropriate epigenetic modifications to ensure the successful transition of the KSHV life cycle(42). Consistent with these, we further validated that the coding region of ORF48 is critical for successful lytic replication of KSHV, through maintaining optimal RTA promoter activities.

Our data show that ORF48 protein is required for an optimal KSHV lytic life cycle, consistent with the functions of its homolog in EBV and MHV68. However, the molecular mechanisms through which this occurs remain to be explored(30–32). Previous findings show that ORF48 inhibits cGAS-STING-based IFNβ promoter activity as well as interaction with PPP6C, a binding partner and a negative regulator of STING(29). Excitingly, we also added results showing ORF48-STING interactions in multiple cell lines. Thus, we originally hypothesized that ORF48 forms a complex with STING and PPP6C and blocks the cGAS/STING pathways to facilitate viral lytic replications. However, we did not observe the enhancement of IFNβ when ORF48 is removed during KSHV lytic infection in any of our iSLK models. This suggests that KSHV might utilize alternative mechanisms to compensate for the ORF48-mediated IFNβ repression, as redundancy is a common strategy employed by pathogens. Indeed, after eliminating redundancy in a KSHV negative background, the removal of ORF48 enhanced the cGAS-STING signaling in our HUVEC-ORF48 cells.

Loss of ORF48 failed to induce STING-dependent IFNβ signaling in the presence of other KSHV genes during lytic reactivation. Although ORF48 is dispensable for IFNβ suppression in this system, loss of ORF48 might still increase the burden for other IFNβ viral inhibitors. Additional mutations on other KSHV ORFs might eventually reach the compensation capacity of KSHV, which compromises KSHV lytic infection. The collective data from these sets of experiments suggest 1. The integrity of ORF48 gDNA and mRNA both contribute to optimal KSHV lytic reactivation 2. ORF48, along with other redundant KSHV genes repress the cGAS/STING pathway during the KSHV lifecycle. 3. ORF48 facilitates KSHV lytic replication through additional unknown mechanisms. The fact that multiple KSHV inhibitors of the cGAS/STING pathways were identified by others and us highlights the critical role of this pathway in counteracting KSHV infection. Further characterization of mechanistic investigation of newly identified IFNβ inhibitors will shed light on KSHV immune evasion, a critical component of KSHV cancer establishment.

## ACKNOWLEDGMENTS

We are grateful to Dr. Blossom Damania for providing the vIL6 monoclonal antibody and immortalized HUVEC. We thank Dr. Rolf Renne for providing plasmid pEPKan-S, E. coli strain GS1783 carrying BAC16, the iSLK.RTA, iSLK.BAC16 and iSLK.219 cells. We thank Mrs. Savannah Hardiman from Dr. Rolf Renne’s lab for providing technical support for the construction of BACmids. We thank Dr. Zsolt Toth for providing RTA promoter-luciferase and RTA plasmids. We thank members of the Ma laboratory for critical readings of the manuscript and helpful discussions. This manuscript is supported by NCI 4R00CA230178, the American Cancer Society Institutional Research Grant, the Department of Molecular Genetics and Microbiology startup funding at UF, and the UF Health Cancer Center startup funding.

